# Super-resolution fluorescence imaging of cryosamples does not limit achievable resolution in cryoEM

**DOI:** 10.1101/2023.03.08.531682

**Authors:** Mart G. F. Last, Willem E.M. Noteborn, Lenard M. Voortman, Thomas H. Sharp

**Affiliations:** Department of Cell and Chemical Biology, Leiden University Medical Centre, 2300 RC Leiden, The Netherlands; Netherlands Centre for Electron Nanoscopy, Leiden University, 2333 AL, Leiden, The Netherlands

## Abstract

Correlated super-resolution cryo-fluorescence and cryo-electron microscopy (cryoEM) has been gaining popularity as a method to investigate biological samples with high resolution and specificity. A concern in this combined method (called SR-cryoCLEM), however, is whether and how fluorescence imaging prior to cryoEM acquisition is detrimental to sample integrity. In this report, we investigated the effect of high-dose laser light irradiation on apoferritin samples prepared for cryoEM with excitation wavelengths commonly used in fluorescence microscopy, and comparing these samples to controls that were kept in the dark. We found that laser illumination, of equal duration and intensity as used in super-resolution cryomicroscopy and in the presence of high concentrations of fluorescent protein, did not affect the achievable resolution in cryoEM, with final reconstructions reaching resolutions of ~1.8 Å regardless of the illumination conditions. The finding that super-resolution fluorescence imaging of cryosamples prior to cryoEM data acquisition does not limit the achievable resolution suggests that super-resolution cryo-fluorescence microscopy and in situ structural biology using cryoEM are entirely compatible.

**Graphical abstract:** 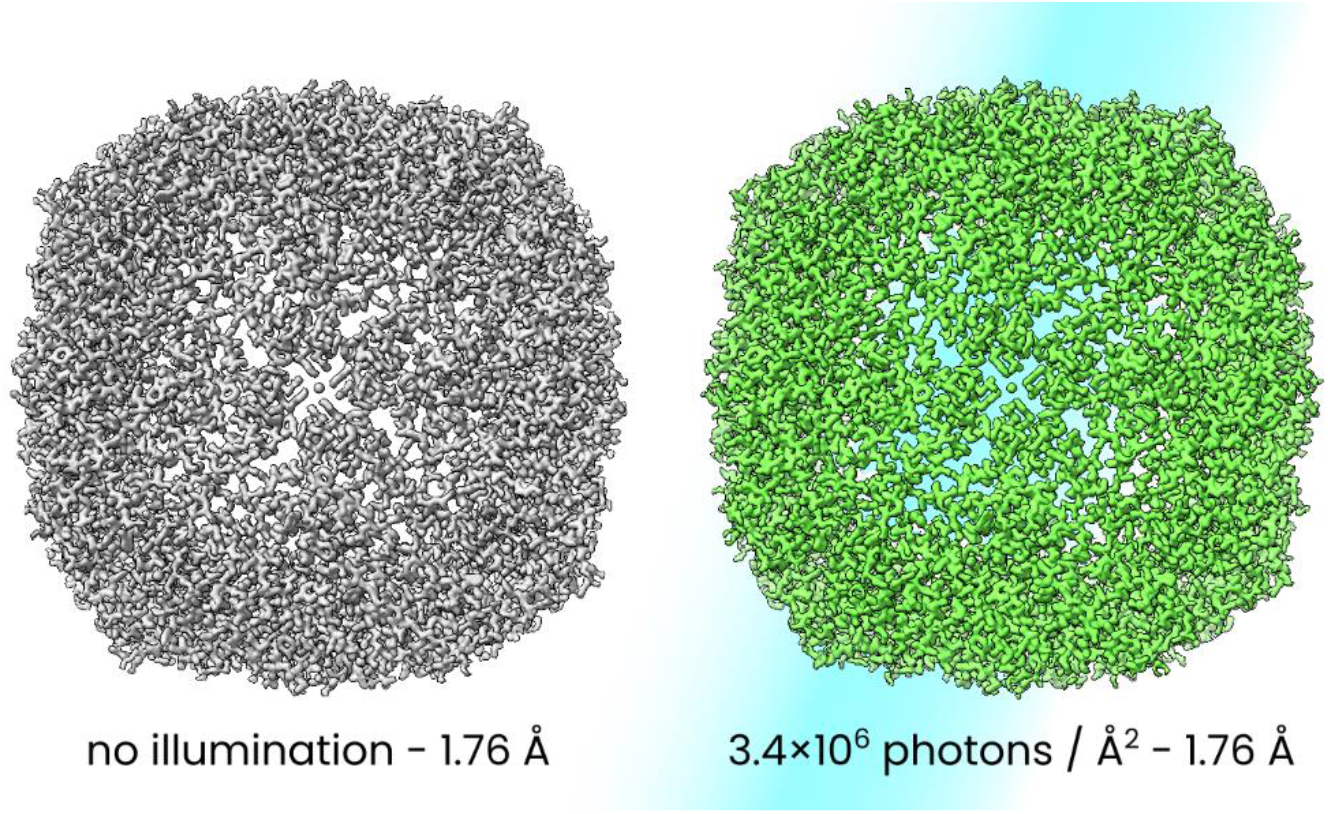

## Introduction

Recent reports demonstrate that by avoiding optical heating of samples in correlated fluorescence and transmission electron cryo-microscopy, high illumination power densities can be used that enable better super-resolution fluorescence imaging^1,2^. While this rules out devitrification as a mode of damaging the sample, it is not clear whether fluorescence inspection of a sample prior to cryoEM imaging damages the sample via other possible mechanisms, such as via the illumination of highly absorptive fluorescent proteins (FPs) with intense laser light. FPs are known to generate reactive oxygen species upon excitation in live-cell fluorescence microscopy^3,4^, to an extent that can lead to cell death^5^. It is currently not known whether this phenomenon could also affect the achievable resolution in cryoEM, but since the reactive species generated by illumination of FPs^4^ and various endogenous chemicals such as flavins^6,7^ are the same as those that have been postulated to account for damage that occurs during cryoEM imaging^8^, it is important to explore this possibility.

In this paper, we investigate the effect of high-power laser illumination on cryosamples by determining the structure of apoferritin, a protein commonly used to benchmark resolution in cryoEM^9^, after exposing samples to intense laser irradiation (up to 3.9 × 10^6^ photons/Å^2^ or 13.8 kJ/cm^2^). By comparing conditions where samples were either irradiated or kept dark, and samples with or without high concentrations of fluorescent protein, we investigate whether light-induced damage to cryosamples occurs under conditions similar to those used in super-resolution fluorescence cryo-microscopy.

## Results

### 1.8 Å structure of apoferritin after laser illumination

In order to test whether intense laser illumination affects the achievable resolution in cryoEM, we performed single-particle analysis on apoferritin samples that were either exposed to intense laser light or kept in the dark as a control. To avoid inconsistencies between test conditions due to differences between samples, we used separate regions on the same grid to test different conditions, i.e., on one grid, we used a number of grid squares for 488 nm illumination, another, distant, region to test the effect of 405 nm illumination, and a third non-adjacent area of the sample for a dark control. Nevertheless, different regions on one grid can still differ in certain parameters, such as the number of suitable holes per grid square. And since the total number of particles in the final reconstruction has a direct influence on the achieved resolution, we limited the total number of particles that were used to generate the final reconstruction to approximately 10^5^, which was the number of particles that remained in the smallest dataset after classification.

Samples were illuminated in a custom-built fluorescence cryo-microscope that we have previously used to determine safe illumination power densities for fluorescence microscopy^2^. Illumination power densities were limited in order to avoid devitrification or melting, yet high enough (100-140 W/cm^2^) to be suitable for single-molecule localization microscopy^2,10–12^. With the knowledge that these photon dose rates are sufficiently low to maintain sample integrity, we now wanted to investigate the effect of the total photon dose.

We initially tested four conditions: exposure to 405 nm, 488 nm, and 561 nm light, plus a non-irradiated control. The illumination power densities that were used were 100, 138, and 138 W/cm^2^, respectively. Individual grid squares were exposed, one at a time, for 100 seconds, corresponding to total doses of 2.0, 3.4, and 3.9 × 10^6^ photons/Å^2^ (see Table 1); for reference, in a typical single-molecule localization microscopy (SMLM) acquisition using rsEGFP2^13^, a widely used reversely switchable fluorescent protein that is also one of the current best labels for cryo-SMLM^12^, we use a dose of 0.2 × 10^6^ 405 nm photons/Å^2^ and 3.4 × 10^6^ 488 nm photons/Å^2^. The 488 nm dose used here is thus typical, while the 405 nm dose is one order of magnitude higher than that used for SMLM image acquisition.

**Table 1.**
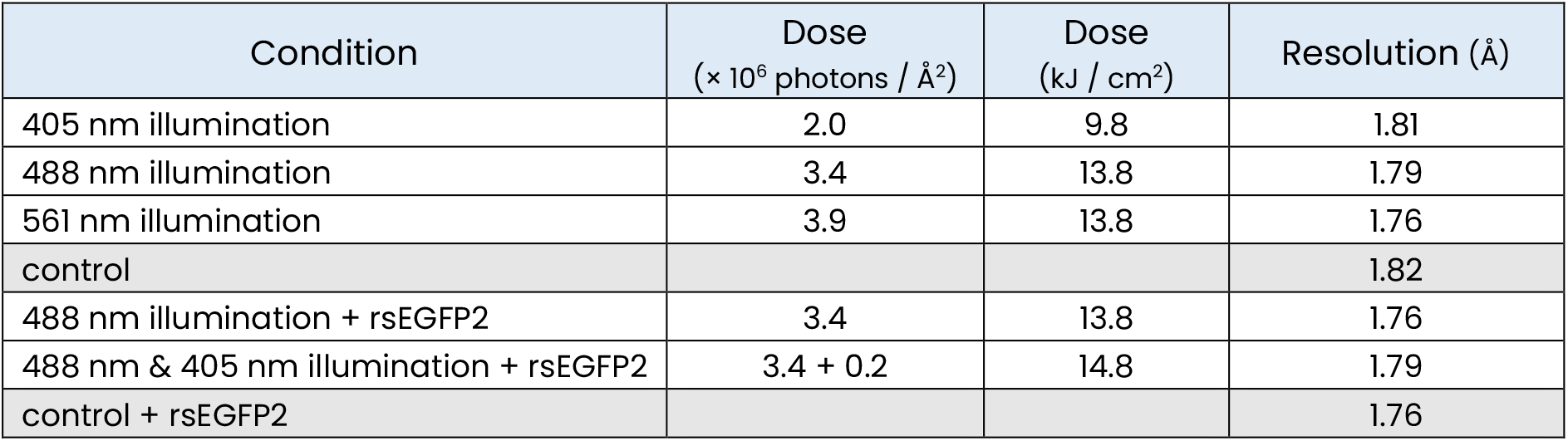
summary of the illumination conditions and resolutions of the final apoferritin structures. The number of particles per dataset was limited to 97000, which was the number found in the dataset with the lowest number of suitable particles (488 nm illumination, see SI for a full overview of dataset and EM parameters). Resolutions are better than or slightly worse than corresponding controls to within 0.03 Å, i.e., illumination does not degrade resolution. Structural differences were also not observed (see Figure 2)

After illumination, we acquired single-particle analysis datasets on the same sample using a 300 kV cryo-transmission electron microscope and processed the resulting data using RELION^14^, which yielded maps at resolutions of 1.81, 1.79, 1.76, and 1.82 Å, for the 405, 488, 561 nm, and dark conditions, respectively (Table 1, Supplementary Table 1). These resolutions are comparable and sufficient to allow the observation of any potential structural defects caused by irradiation^15^.

### The presence of fluorescent protein does not affect resolution

Next, we performed similar experiments with samples that contained a high concentration of rsEGFP2 alongside apoferritin (Figure 1). By itself, apoferritin does not have a high extinction coefficient at any visible wavelength, and we therefore posited that the addition of rsEGFP2 would thus lead to increased absorption of light, as well as subject apoferritin to the effect of reactive species generated by rsEGFP2 upon excitation, thereby rendering conditions comparable to those that occur in samples for fluorescence microscopy.

**Figure 1.**
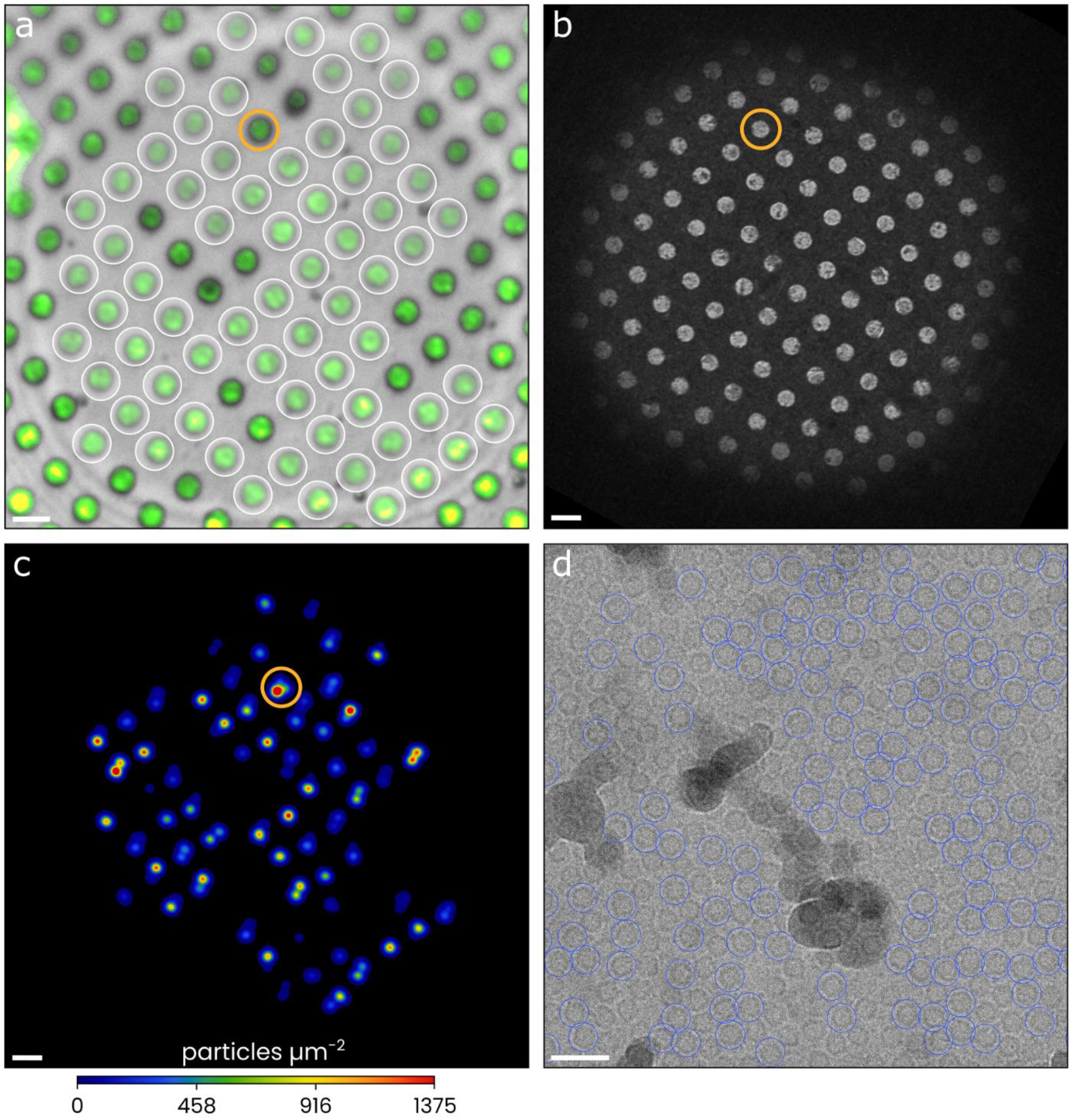
representative light and electron microscopy images of the samples used in this study. a) A cryo-light microscopy image of a grid square, showing rsEGFP2 fluorescence (green) overlayed on a reflected light image (grey). CryoEM micrographs were acquired in the holes encircled in white. b) A cryoEM overview image of the same region of the sample as in panel a. c) A heat map of the spatial distribution of particles that were used in the final reconstruction. d) A micrograph acquired in the hole indicated (in orange) in the adjacent images. The micrograph displays numerous apoferritin molecules, a fraction (blue circles) of which was used for the final reconstruction. Scale bars: 2 μm (a, b, c), 30 nm (d).

To validate that protein structure is not impacted by conditions encountered in a super-resolution imaging experiment, we illuminated this sample using the same acquisition routine that we employ in SMLM: a sequence of ten fluorescence images (10 × 488 nm × 100 ms at 138 W/cm^2^) followed by a near-UV activation pulse (1 × 405 nm × 100 ms at 100 W/cm^2^), and a reflected light image (1 × 528 nm × 100 ms at less than 1 W/cm^2^), repeated 100 times (Figure 1a). As a control, we also acquired cryoEM datasets for regions of the sample exposed to just 488 nm illumination, as well as a non-illuminated control. This resulted in three additional maps with a resolution of 1.76, 1.79, and 1.76 Å (Table 1).

### Super-resolution imaging does not limit achievable cryoEM resolution

To assess whether any structural changes occurred to apoferritin as a consequence of the high-intensity laser illumination, we fitted molecular models into each of the seven density maps and compared the resulting structures (Figure 2, Supplementary Figures 1-2). The overall degree of similarity was high, with only 0.02 Å root mean square deviation between the position of alpha carbon atoms of the seven models. No notable differences between the models were observed at the single-residue level, with the exception of a number of solvent-facing residues (Ser4, His57, Glu162) which were fitted in different rotamer states in regions where the density maps were ambiguous. Inspection of the density maps also did not reveal any marked differences. In conclusion, laser illumination of cryosamples with dose rates comparable to and total photon doses equal to or higher than those used in super-resolution fluorescence microscope did not have an effect on the achievable resolution, nor on the final structural models.

**Figure 2.**
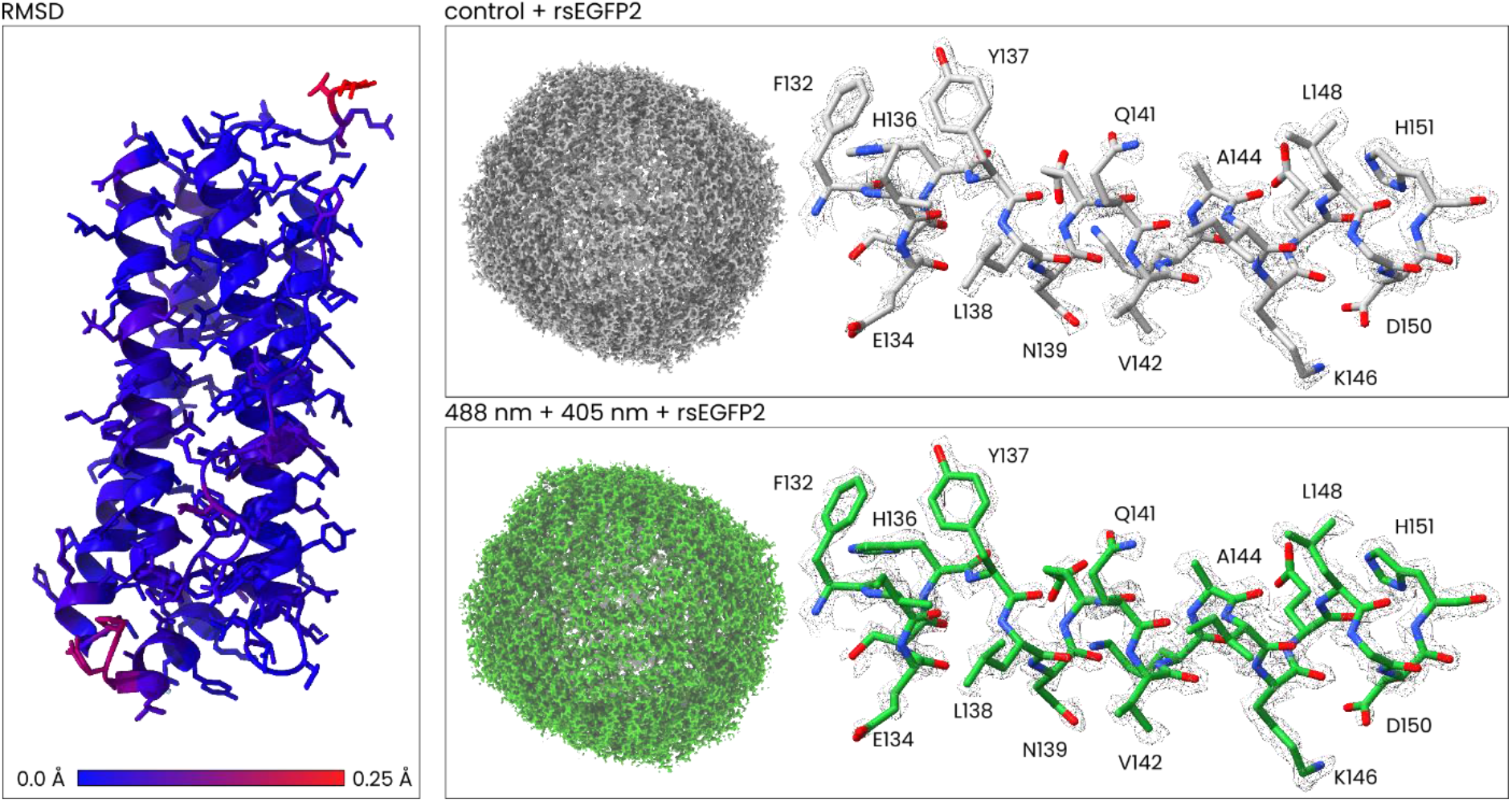
Structures of apoferritin after no exposure and after illumination equivalent to that used in a super-resolution fluorescence acquisition. Images of the seven full maps and regions showing the map and model fit can be found in the supplementary information. Here, we compare the maps obtained for the 488 nm + 405 nm illumination in presence of rsEGFP2 with a dark control. Left: an overlay of the two models (showing a single monomer), with residues coloured in proportion to the root mean square deviation between the position of the alpha carbon atoms of the two models. Right: full maps and detailed views of residues 132 to 151 for the control (top) and laser illuminated datasets (bottom). A number of residues are indicated, including some (e.g. E134, Y137, H151) that would have been most demonstrative of damage^16^, should any have occurred.

## Discussion

In this report, we have presented data that addresses the question of whether exposure to intense light sources damages cryoEM samples. This is an important open question for the prospect of combining (super-resolution) fluorescence microscopy with cryoEM – a novel methodology that has been gaining popularity in recent years^17,18^. The correlation of fluorescence and electron microscopy promises a number of highly advantageous applications in structural cell biology, such as accurate targeting of cryoEM data acquisition via localization of fluorescently labelled structures of interest and the possibility of identifying densities observed in cryo-electron tomographic volumes by correlation with single molecule localization microscopy particle maps. However, whether fluorescence imaging of cryosamples is detrimental to the achievable resolution had not yet been addressed. Our findings conclusively show that high-power laser illumination of cryosamples does not lead to a reduction in the achievable resolution, indicating that the potential advantages offered by combining super-resolution fluorescence and cryoEM imaging need not come at the cost of resolution.

With apoferritin as a representative non-photosensitive protein in the presence of the photosensitizing fluorophore rsEGFP2, we expect that our findings are also valid for cellular samples containing fluorescent fusion proteins. Given that there are no observable differences in apoferritin at <2.0 Å, we expect it to be rare for fluorescence imaging of cryosamples to lead to any observable differences in *in situ* cryo-electron tomography datasets, where resolution often does not reach this level. Indeed, a recent report^19^ demonstrated that brief illumination with 185 – 280 nm UVC light had no effect on the obtainable resolution in cryo-electron tomography above 22.6 Å, or in single-particle analysis above 2.1 Å; however, the total doses used in that report were almost 1,000,000× lower than those used here.

In conclusion, we have now shown that irradiating cryosamples with high dose rates and high total doses of visible light does not have a measurable effect on the achievable resolution of cryoEM. This is an important validation of the applicability of super-resolution fluorescence localization of structures of interest within a cryoEM imaging workflow: it shows that super-resolution fluorescence imaging is entirely compatible with cryoEM.

## Methods

### Sample preparation

Samples were prepared by glow discharging 1.2/1.3 300 mesh UltrAuFoil (QuantiFoil Micro Tools GmbH, Großlöbichau, Germany) holey gold grids^20^, depositing 3.5 μL of either 4 mg/mL (8.9 μM) apoferritin or a mixture containing 3.6 mg/mL (8 μM) apoferritin and 30 μM rsEGFP2 onto the grids, blotting for 3 seconds, and plunge freezing in liquid ethane using a Vitrobot Mark IV (Thermo Fisher Scientific) with the environmental chamber set to 22°C and 100% humidity.

### Laser illumination of cryosamples

After plunge freezing, samples were irradiated in a custom-built fluorescence cryo-microscope^2^ that is equipped with high powered 405, 488, and 561 nm lasers (100-150 mW, LightHUB, Omicron-Laserage GmbH, Rodgau, Germany). The illumination power density was determined using a S130C power meter (Thorlabs, Newton, NJ, USA) to measure the light output in the focal plane at the front aperture of the objective lens and dividing that value by the size of the illuminated spot, as measured directly from images. The illuminated field size was restricted to a circle of approximately 86 μm in diameter by use of a field stop in order to ensure that only one grid square would be illuminated at a time.

Samples were mounted in a CMS196v3 liquid nitrogen cooled light microscopy stage (Linkam Scientific Instruments Ltd., Salfords, UK) and videos (1000 frames with 100 ms exposure, for a 100 second total exposure time) were acquired in similar fashion to how we acquire data for super-resolution fluorescence, using the following illumination power densities: 138 W/cm^2^ for the 488 nm laser, 100 W/cm^2^ for 405 nm, and 138 W/cm^2^ for 561 nm, corresponding to a total of, respectively, 3.4, 2.0, and 3.9 × 10^6^ photons/Å^2^.

For the purpose of correlation with cryoEM overview images, reflected light images were acquired using a low-powered (< 1 W/cm^2^) 528 nm LED light source for illumination.

The 561 nm illumination dataset was acquired on a different grid than the 405 nm and 488 nm datasets and their respective control dataset - i.e., no separate control was acquired for the 561 nm illumination dataset. Since the final resolution under this condition reached 1.75 Å, a value similar to that of both the earlier controls (with and without rsEGFP2), we decided against acquisition of a third, separate, control dataset for the 561 nm condition.

### CryoEM data acquisition

CryoEM data acquisition was performed at the Netherlands Centre for Electron Nanoscopy (NeCEN) on a Titan Krios microscope operating at 300 kV (Thermo Fisher Scientific, Eindhoven, The Netherlands) equipped with a BioQuantum energy filter using a slit width of 20 eV and K3 direct electron detector (Gatan, Inc, Pleasanton, CA, USA). Movies were collected using EPU software (Thermo Fisher Scientific) in counted super-resolution mode, using aberration free image shift (AFIS) at a nominal magnification of 130.000 (0.328 Å/pixel). The defocus range was set from −0.5 to −1.5 μm and the total accumulated dose to 50 electrons/Å^2^. Further details per dataset are reported in Supplementary Table 1.

### CryoEM data processing

CryoEM data processing was performed in RELION-4.0^14^. Super-resolution movies were corrected for beam-induced drift and binned to a pixel size of 0.656 Å/pixel using MotionCor2^21^ and CTF estimation was done using CTFFIND-4.1.18^22^. Particle picking was done using the standard Topaz neural network^23^ and after extraction particles were submitted to 2D classification. 2D classes were ranked using the RELION class ranker^14^. Classes with a score of 0.5 and higher were selected for 3D refinement. Ab-initio models were generated using octahedral symmetry. Using the same symmetry, 3D reconstructions were obtained after 3D refinement, CTF refinement (beam-tilt and aberration correction, anisotropic magnification estimation, and per-particle defocus and per-micrograph astigmatism correction), Bayesian polishing, a second round of CTF refinement and a last round of 3D refinement. Final reconstructions were sharpened and locally filtered using RELION post-processing.

### Model building

Using a recently determined apoferritin structure^24^ (protein database (PDB) code 6Z6U) as a starting model, we first manually added magnesium atoms at known coordination sites and removed all water molecules using University of California San Francisco (UCSF) ChimeraX^25^ and ISOLDE^26^. Then, the full model (all 24 monomers) fit was automatically refined using Phenix real-space refine^27^, after which water molecules were added back in the model using phenix.douse followed by a second, final refinement of the model. Accession codes for the final models as well as maps are listed in Supplementary Table 1.

### Mapping particle positions into cryoEM overview images

In order to generate heat maps of the spatial distribution of the particles that made it in to the final reconstructions (e.g. Figure 1c) and overlay these onto overview images of a grid square, we had to convert the pixel coordinates of particles within micrographs into a corresponding cryoEM stage position (the stage position is known for overview images). This involves a number of calibrations: 1) converting a micrograph’s beam shift, which is stored in terms of arbitrary units, into spatial units, 2) finding the angle of rotation between the beam shift coordinate system and the stage coordinate system, and 3) finding the angle of rotation between the stage coordinate system and the image coordinate system. Lacking access to the calibration registry of the TEM used for acquisition, we used an approximated calibration, due to which the accuracy of the colocalization in Figure 1c is limited with ~0.0-1.0 μm error. This error is less than the spacing between holes (1.3 μm), so that one can still visually pair holes and localizations. The heat map was blurred somewhat in order to aid visual inspection; localization clouds for one micrograph originally would not exceed the size of the micrograph, which was 390 nm × 270 nm.

## Acknowledgements

We thank A. Moutsiopoulou and M. Lamers for providing us with purified apoferritin. This research was supported by the following grants to THS: European Research Council Grant 759517; The Netherlands Organization for Scientific Research Grants OCENW.KLEIN.291 and VI.Vidi.193.014, and by the Netherlands Electron Microscopy Infrastructure (NEMI), project number 184.034.014 of the National Roadmap for Large-Scale Research Infrastructure of the Dutch Research Council (NWO).

## References

1 Dahlberg, P. D., Perez, D., Hecksel, C. W., Chiu, W. & Moerner, W. E. Metallic support films reduce optical heating in cryogenic correlative light and electron tomography. Journal of Structural Biology 214, 107901 (2022).

2 Last, M. G. F., Tuijtel, M. W., Voortman, L. M. & Sharp, T. H. Selecting optimal support grids for super-resolution cryogenic correlated light and electron microscopy. bioRxiv (2022).

3 Icha, J., Weber, M., Waters, J. C. & Norden, C. Phototoxicity in live fluorescence microscopy, and how to avoid it. Bioessays 39 (2017). https://doi.org:10.1002/bies.201700003

4 Trewin, A. J. et al. Light-induced oxidant production by fluorescent proteins. Free Radic Biol Med 128, 157–164 (2018). https://doi.org:10.1016/j.freeradbiomed.2018.02.002

5 Pletneva, N. V. et al. Crystal Structure of Phototoxic Orange Fluorescent Proteins with a Tryptophan-Based Chromophore. PLOS ONE 10, e0145740 (2015). https://doi.org:10.1371/journal.pone.0145740

6 Eichler, M., Lavi, R., Shainberg, A. & Lubart, R. Flavins are source of visible-light-induced free radical formation in cells. Lasers in Surgery and Medicine 37, 314–319 (2005). https://doi.org/10.1002/lsm.20239

7 Baumler, W., Regensburger, J., Knak, A., Felgentrager, A. & Maisch, T. UVA and endogenous photosensitizers--the detection of singlet oxygen by its luminescence. Photochem Photobiol Sci 11, 107–117 (2012). https://doi.org:10.1039/c1pp05142c

8 Naydenova, K. et al. On the reduction in the effects of radiation damage to two dimensional crystals of organic and biological molecules at liquid-helium temperature. Ultramicroscopy 237, 113512 (2022). https://doi.org:10.1016/j.ultramic.2022.113512

9 Danev, R., Yanagisawa, H. & Kikkawa, M. Cryo-EM performance testing of hardware and data acquisition strategies. Microscopy 70, 487–497 (2021). https://doi.org:10.1093/jmicro/dfab016

10 Dahlberg, P. D. et al. Cryogenic single-molecule fluorescence annotations for electron tomography reveal in situ organization of key proteins in Caulobacter. Proceedings of the National Academy of Sciences 117, 13937–13944 (2020). https://doi.org:10.1073/pnas.2001849117

11 Moser, F. et al. Cryo-SOFI enabling low-dose super-resolution correlative light and electron cryo-microscopy. Proc Natl Acad Sci U S A 116, 4804–4809 (2019). https://doi.org:10.1073/pnas.1810690116

12 Tuijtel, M. W., Koster, A. J., Jakobs, S., Faas, F. G. A. & Sharp, T. H. Correlative cryo super-resolution light and electron microscopy on mammalian cells using fluorescent proteins. Scientific Reports 9, 1369 (2019). https://doi.org:10.1038/s41598-018-37728-8

13 Grotjohann, T. et al. rsEGFP2 enables fast RESOLFT nanoscopy of living cells. Elife 1, e00248 (2012). https://doi.org:10.7554/eLife.00248

14 Kimanius, D., Dong, L., Sharov, G., Nakane, T. & Scheres, S. H. W. New tools for automated cryo-EM single-particle analysis in RELION-4.0. Biochem J 478, 4169–4185 (2021). https://doi.org:10.1042/BCJ20210708

15 Hattne, J. et al. Analysis of Global and Site-Specific Radiation Damage in Cryo-EM. Structure 26, 759–766.e754 (2018). https://doi.org/10.1016/j.str.2018.03.021

16 Matthies, D., Bartesaghi, A., Merk, A., Banerjee, S. & Subramaniam, S. Residue Specific Radiation Damage of Protein Structures using High-Resolution Cryo-Electron Microscopy. Biophysical Journal 108, 190a (2015). https://doi.org:10.1016/j.bpj.2014.11.1052

17 Dahlberg, P. D. & Moerner, W. E. Cryogenic Super-Resolution Fluorescence and Electron Microscopy Correlated at the Nanoscale. Annu Rev Phys Chem 72, 253–278 (2021). https://doi.org:10.1146/annurev-physchem-090319-051546

18 DeRosier, D. J. Where in the cell is my protein? Quarterly Reviews of Biophysics 54, e9 (2021). https://doi.org:10.1017/S003358352100007X

19 Depelteau, J. S. et al. UVC inactivation of pathogenic samples suitable for cryo-EM analysis. Communications Biology 5, 29 (2022). https://doi.org:10.1038/s42003-021-02962-w

20 Russo, C. J. & Passmore, L. A. Ultrastable gold substrates for electron cryomicroscopy. Science 346, 1377–1380 (2014). https://doi.org:10.1126/science.1259530

21 Zheng, S. Q. et al. MotionCor2: anisotropic correction of beam-induced motion for improved cryo-electron microscopy. Nat Methods 14, 331–332 (2017). https://doi.org:10.1038/nmeth.4193

22 Rohou, A. & Grigorieff, N. CTFFIND4: Fast and accurate defocus estimation from electron micrographs. J Struct Biol 192, 216–221 (2015). https://doi.org:10.1016/j.jsb.2015.08.008

23 Bepler, T. et al. Positive-unlabeled convolutional neural networks for particle picking in cryo-electron micrographs. Nat Methods 16, 1153–1160 (2019). https://doi.org:10.1038/s41592-019-0575-8

24 Yip, K. M., Fischer, N., Paknia, E., Chari, A. & Stark, H. Atomic-resolution protein structure determination by cryo-EM. Nature 587, 157–161 (2020). https://doi.org:10.1038/s41586-020-2833-4

25 Pettersen, E. F. et al. UCSF ChimeraX: Structure visualization for researchers, educators, and developers. Protein Sci 30, 70–82 (2021). https://doi.org:10.1002/pro.3943

26 Croll, T. I. ISOLDE: a physically realistic environment for model building into low-resolution electron-density maps. Acta Crystallogr D Struct Biol 74, 519–530 (2018). https://doi.org:10.1107/S2059798318002425

27 Afonine, P. V. et al. Real-space refinement in PHENIX for cryo-EM and crystallography. Acta Crystallogr D Struct Biol 74, 531–544 (2018). https://doi.org:10.1107/S2059798318006551

