## Supplementary Information for "Super-resolution fluorescence imaging of cryosamples does not limit achievable resolution in cryoEM"

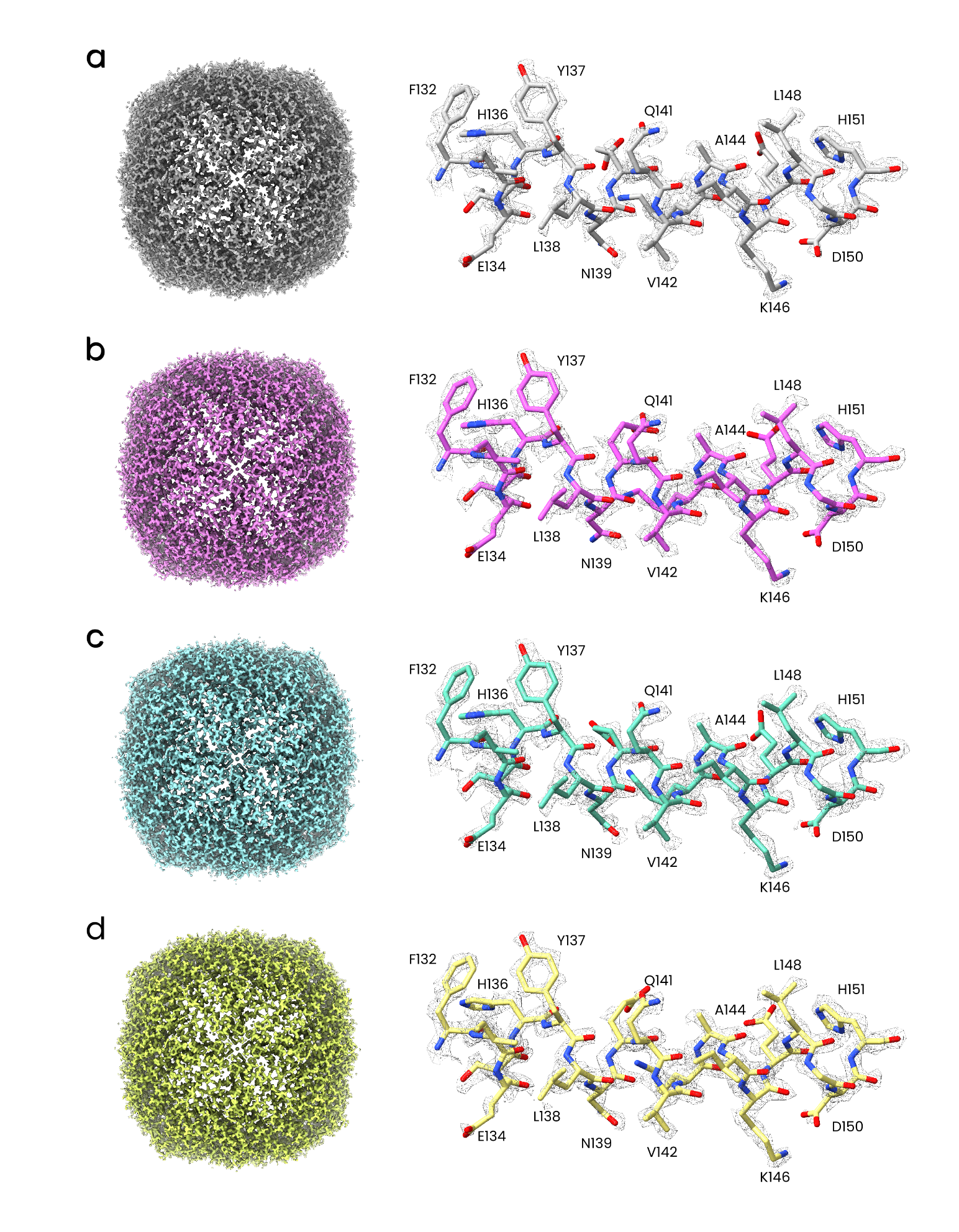

Supplementary Figure 1 – Maps and models of apoferritin for the (**a**) dark, (**b**) 405 nm, (**c**) 488 nm, and (**d**) 561 nm illuminated, no rsEGFP2 conditions (respectively, top to bottom). The full density map, at a threshold of 0.025, is shown to the left. On the right, the map and model for residues 132-151 is displayed, with a number of the residues labelled.

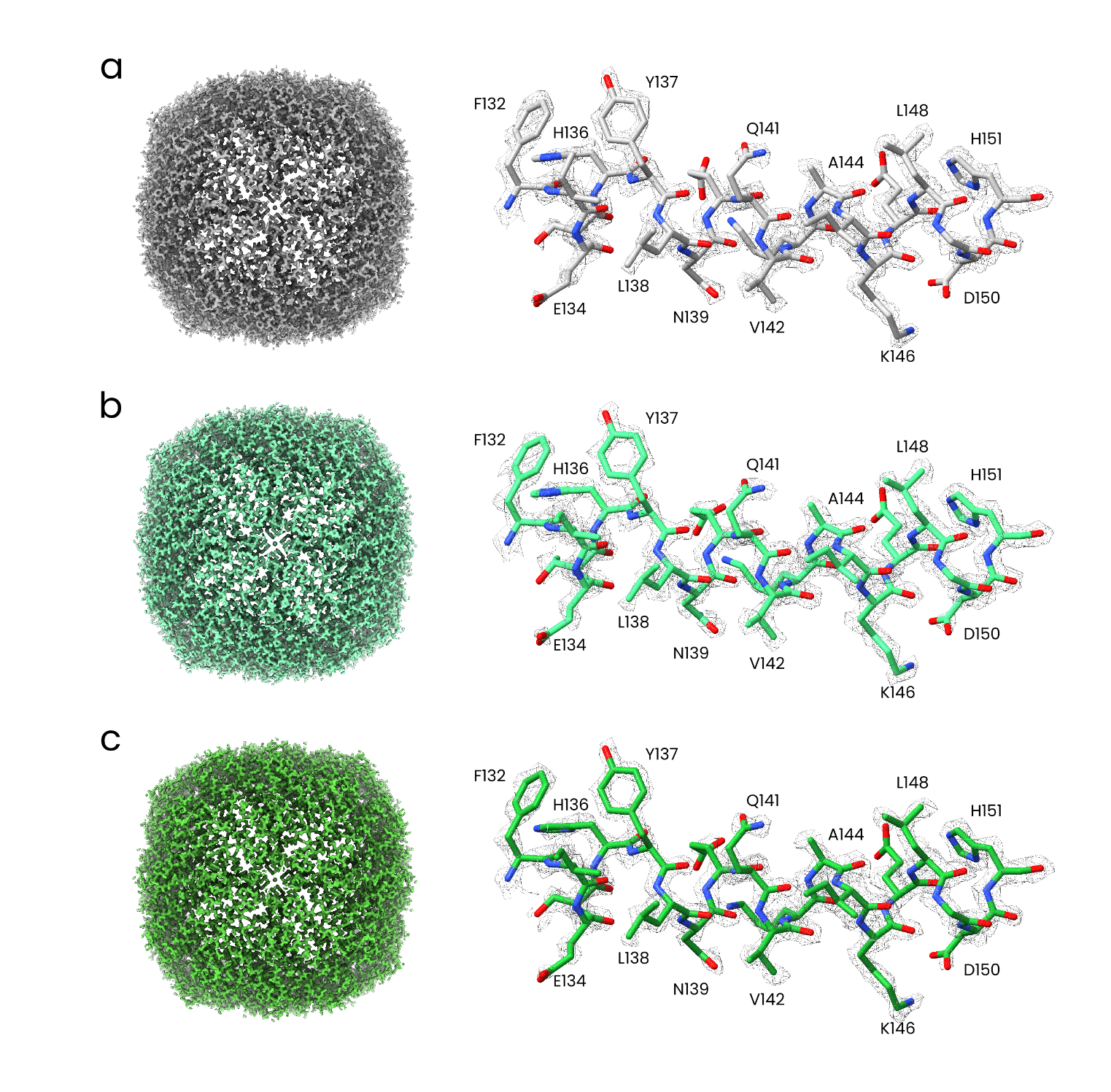

Supplementary Figure 2 – Maps and models of apoferritin for the (**a**) dark, (**b**) 488 nm, and (**c)** 405 nm + 488 nm illuminated and with rsEGFP2 added conditions. The full density map, at a threshold of 0.025, is shown to the left. On the right, the map and model for residues 132-151 is displayed, with a number of the residues labelled.

| Condition: | Dark | 405 nm | 488 nm | 561 nm | Dark + rsEGFP2 | 488 nm + rsEGFP2 | 488 nm + 405 nm + rsEGFP2 |
| --- | --- | --- | --- | --- | --- | --- | --- |
| EM and refinement parameters | | | | | | | |
| No. of micrographs | 1612 | 1688 | 1825 | 1938 | 1440 | 1336 | 1610 |
| No. of particles total | 2043237 | 2036041 | 2317586 | 2501778 | 1677002 | 1675076 | 1904135 |
| No. of particles / micrograph | 1268 | 1206 | 1270 | 1291 | 1165 | 1254 | 1183 |
| EMBD accession code | EMD-16784 | EMD-16783 | EMD-16785 | EMD-16786 | EMD-16787 | EMD-16789 | EMD-16788 |
| PDB accession code | 8CPS | 8CPM | 8CPT | 8CPU | 8CPV | 8CPX | 8CPW |
| No. of final particles | 170782 | 144851 | 98688 | 246513 | 228943 | 177864 | 126701 |
| Restricted no. of final particles | 97000 | 97000 | 97000 | 97000 | 97000 | 97000 | 97000 |
| Map resolution  (Å, FSC = 0.143 threshold) | 1.82 | 1.81 | 1.79 | 1.76 | 1.76 | 1.76 | 1.79 |
| Map sharpening B factor (Å^2^) | -44 | -43 | -39 | -40 | -43 | -42 | -43 |
| Model composition | | | | | | | |
| Protein chains (#) | 24 | 24 | 24 | 24 | 24 | 24 | 24 |
| Nonhydrogen atoms (#) | 36034 | 36090 | 36024 | 35823 | 35855 | 35827 | 36004 |
| Residues (#) | 4152 | 4152 | 4152 | 4152 | 4152 | 4152 | 4152 |
| Water (#) | 1916 | 1972 | 1906 | 1705 | 1737 | 1709 | 1886 |
| Ligands (#) | NA:24 MG: 14 | NA: 24  MG: 14 | NA: 24  MG: 14 | NA: 24  MG: 14 | NA: 24  MG: 14 | NA: 24  MG: 14 | NA: 24  MG: 14 |
| Bond lengths (Å, # > 4 σ) | 0.003 (0) | 0.005 (0) | 0.004 (0) | 0.004 (0) | 0.006 (0) | 0.005 (0) | 0.005 (0) |
| Bond angles (°, # > 4 σ) | 0.659 (0) | 0.847 (0) | 0.734 (0) | 0.717 (0) | 0.924 (1) | 0.812 (0) | 0.846 (0) |
| MolProbity score | 1.07 | 1.41 | 0.84 | 1.26 | 1.04 | 1.29 | 1.13 |
| Clash score | 2.82 | 7.50 | 1.25 | 4.95 | 2.58 | 5.37 | 3.39 |
| Ramachandran plot  Outliers % | 0.00 | 0.00 | 0.00 | 0.00 | 0.00 | 0.00 | 0.00 |
| Ramachandran plot  Allowed % | 1.04 | 1.04 | 1.05 | 0.99 | 1.04 | 1.04 | 1.04 |
| Ramachandran plot  Favored % | 98.96 | 98.96 | 98.96 | 99.01 | 98.86 | 98.96 | 98.96 |
| Model resolution  (Å, FSC = 0.5 threshold) | 1.8 | 1.9 | 1.8 | 1.8 | 1.8 | 1.8 | 1.8 |

Supplementary Table 1 – SPA and model parameters plus accession codes for each of the maps and models.
